# Awake infant fMRI: Insights from more than 750 scanning sessions

**DOI:** 10.1101/2025.02.20.636736

**Authors:** Lillian Behm, Tristan S. Yates, Juliana E. Trach, Dawoon Choi, Haoyu Du, Camille Osumah, Ben Deen, Heather L. Kosakowski, Emily M. Chen, Frederik S. Kamps, Halie A. Olson, Cameron T. Ellis, Rebecca Saxe, Nicholas B. Turk-Browne

## Abstract

Functional magnetic resonance imaging (fMRI) in awake infants has the potential to reveal how the early developing brain gives rise to cognition and behavior. However, awake infant fMRI poses significant methodological challenges that have hampered wider adoption. The present work takes stock after the collection of a substantial amount of awake infant fMRI data across multiple studies from two labs at different institutions. These data were leveraged to glean insights on participant recruitment, experimental design, and data acquisition that could be useful to consider for future studies. Across 766 fMRI sessions with awake infants aged 1–36 months, the authors explored the factors that influenced how much usable data were obtained per session. The age of an infant predicted whether they would successfully enter the scanner (younger more likely) and, if they did enter, the number of minutes of functional data collected (linear, younger more) and retained after preprocessing with lab-specific protocols or harmonized motion exclusion thresholds (quadratic, 12–24 months more than younger and older). The amount of functional data retained was also influenced by assigned sex (female more), experimental paradigm (movies better than blocks and events), and stimulus content (social better than abstract). There were many differences in the research approach between labs making head-to-head comparisons difficult, but Yale was more likely to get infants into the scanner, MIT collected more data from infants who entered, and the amount of data retained after preprocessing did not differ statistically between labs (9 minutes). In addition, the authors assessed the value of attempting to collect multiple experiments per session, an approach that yielded more than one usable experiment averaging across all sessions. Although any given scan is unpredictable, these findings support the feasibility of awake infant fMRI and suggest practices to optimize future research.

## Introduction

The human brain develops more rapidly during infancy than any other period in life (1). This intense neurodevelopment is accompanied by the emergence of many foundational cognitive processes and behaviors (2–4). Until recently, researchers were limited in their ability to determine how changes in infant brain function give rise to cognition.

Scalp-based methods such as electroencephalography (EEG), functional near-infrared spectroscopy (fNIRS), and magnetoencephalography (MEG) have valuable strengths for infant studies but have relatively poor spatial resolution and cannot definitively resolve deep-brain structures (5). Functional magnetic resonance imaging (fMRI) is a complementary tool that can probe brain activity during infancy with improved spatial resolution (6) relative to scalp-based methods. Though once considered to be incompatible for use with infants given a host of methodological constraints, fMRI has now been successfully adapted for both sleeping and awake infant studies and is beginning to provide critical insights into early development (7, 8).

There is considerable fMRI research in sleeping infants, akin to adult resting-state data (9), but much less awake infant fMRI. The earliest study of this type reported that cortical language regions (left superior temporal gyrus and angular gyrus) responded more strongly to forward versus backward speech in both awake and asleep infants, but critically, that dorsolateral prefrontal cortex responded differentially only in awake infants (10). This was the first demonstration that performing fMRI in an awake state can benefit understanding of the neural mechanisms of infant cognition. Indeed, nearly all fMRI studies in older children and adults are performed while they are awake, enabling a wide range of perceptual and cognitive tasks and the ability to link brain activity to concurrent behaviors such as eye movements, memory recall, and decision-making.

Over the past few years, there has been a renaissance of awake infant fMRI studies focusing on the early organization of the visual system. This body of work includes a studies of lower-level retinotopic organization, showing the existence of areas V1-V4 in infants as young as 5 months (11, 12). In the dorsal visual pathway, infants as young as five weeks show responses in putative motion regions (13, 14). In the ventral visual pathway, awake infant fMRI has revealed higher-level category organization, with preferential responses in the fusiform face area (FFA), parahippocampal place area (PPA), and extrastriate body area (EBA) in infants from 2-9 months (15, 16). In one study, FFA responses differed between infants tested before versus after the first COVID-19 pandemic lockdown, suggesting a potential role for experience and environmental factors (17) in the development of the visual system. Overall, these studies show that key aspects of the visual system are surprisingly mature in infancy.

Awake infant fMRI has also been used to study cognitive processes beyond perception. This includes the ability of infants to orient attention through eye movements, the basis of the “looking time” dependent measure that dominates behavioral research on infant cognition (18). In a Posner cuing task, frontoparietal regions of adult attention networks were recruited when infants 3–12 months old reoriented attention to an invalidly cued target location (19). fMRI data have also suggested that infants segment continuous perceptual information into discrete cognitive events. A hidden Markov model applied during a movie-watching task revealed that infants had fewer and longer neural events than adults (20), providing provocative clues into the nature of their experience. Finally, awake infant fMRI has been used to study the hippocampus, a deep-brain region critical for learning and memory not easily accessible to other infant-friendly techniques. The lack of memories from infancy later in life (infantile amnesia) suggested that the hippocampus may have a protracted maturation, yet hippocampal activity in infants has now been shown to track statistical learning of temporal regularities (21) and supports the encoding of memories for single items (22). These studies provide novel insights, and there remain countless open questions that awake infant fMRI could help answer.

Why, then, do relatively few labs use this technique? Although the findings above are a testament to feasibility, awake infant fMRI poses several major challenges. For example, fMRI is prone to motion artifacts, yet infants move incessantly while awake and, unlike older children or adults, cannot comprehend or comply with instructions to remain still during a scan. Additionally, unlike behavioral studies, EEG, and fNIRS, fMRI can require separation of the infant from the parent for placement in the bore (though the parent usually remains nearby). Furthermore, infants have a limited attention span of a few minutes, not nearly enough time to collect the amount of task-based fMRI data typical of adult studies. The combination of these and other challenges make awake infant scans labor-intensive for researchers and greatly limit the amount of usable data collected during a session.

Given the promise of awake infant fMRI, researchers may benefit from a better understanding of how these challenges impact data retention and how they might be mitigated. Significant efforts have been made for fMRI research in other populations — adults, children, and sleeping infants/toddlers — to assess data retention and minimize data loss (23–26). This self-reflection has not yet occurred for awake infant fMRI, likely because of the small number of labs conducting such studies. Early adopters of awake infant fMRI utilize open-science practices to share detailed methods, hardware validation, software packages, datasets, and analysis tools (6, 27–29). Even armed with these resources, researchers hoping to acquire awake infant fMRI data may still wonder about the feasibility and success rates they can expect. The goal of the present work is to address these unknowns and help to shape expectations based on two well-tested methods for awake infant fMRI data collection.

The present work aggregates data from two labs at different institutions who specialize in this technique. Having collected thousands of minutes of usable fMRI data from hundreds of awake infants over the past decade, the authors are in a unique position to share their experiences with participant recruitment, task design, and data acquisition. The current paper describes the protocols that each lab has carefully developed and refined to collect data and conduct research using fMRI with awake infants. Though the authors have collected data from a wide range of ages (1–36 months), the protocols described in this paper were initially developed for participants up to two years old. Indeed, the bulk of the data is from participants under one year old and there is not sufficient data to make definitive claims about toddlers specifically. Therefore, the authors use a broad definition of “infant” to refer to all participants in the current sample.

In the present work, the authors have combined their respective datasets to explore which factors influence scanning success for both groups. The authors investigate factors that predict whether an infant will enter the scanner and, if they do, the amount of functional data obtained. The authors analyze sources of data loss such as head motion and looking behavior. The authors test how scanning success is influenced by the number of repeat sessions, experimental paradigm, and state and trait characteristics of the infant (reported by parents). The authors conclude with a discussion contrasting their approaches and reflecting upon the strengths and challenges of implementing each. They then consider how these results and insights may inform future awake infant fMRI studies. These observations are not meant to be prescriptive or to serve as best practices, but rather to share what has (and has not) worked so far for these groups given their respective facilities and goals. How these insights generalize to other labs remains to be tested, but the authors are hopeful that this work will allow others to benefit from their experiences and inform their approaches.

## Methods

### Yale University

#### Participants

The Turk-Browne Lab at Yale University collected data from 115 unique infants aged 3.3-32.6 months (mean, *M* = 10.91; standard deviation, *SD* = 6.04; 152 female, 11 sex not reported) across a total of 304 scanning sessions. Data collection for the project took place at three academic research-dedicated scanning locations over time: the Scully Center for the Neuroscience of Mind and Behavior at Princeton University from February 2016 to May 2017 (27 sessions), Magnetic Resonance Research Center (MRRC) at Yale University from July 2018 to Dec 2018 (36 sessions), and the Brain Imaging Center (BIC) at Yale University from January 2019 to May 2023 (241 sessions). The infants recruited from the greater Princeton, New Jersey area primarily resided in a predominantly white (Non-Hispanic) upper-middle-class community where English is the primary language spoken. Infants recruited from the greater New Haven, Connecticut area resided in a racially, ethnically, and socioeconomically diverse community where English is the primary language spoken. A parent and/or legal guardian provided informed consent on behalf of their child. This research was approved by the Institutional Review Boards at Princeton University and Yale University.

#### Recruitment and orientation

Recruitment methods differed across locations. At Princeton University, the research was advertised via flyers, engagement in community activities, and word of mouth. At Yale University, recruitment was conducted via the Yale Baby School, a multi-laboratory initiative for community engagement in developmental research. A team member visited families in the Maternity and New-born Unit at the Yale New Haven Hospital in the days immediately following birth to introduce the Yale Baby School and obtain contact information if interested. Of the 3,579 families who provided contact information since January 2018, 722 (20.2%) eventually enrolled in the Yale Baby School. These families were then allocated to one of several approved studies based on eligibility criteria, with the possibility of subsequent re-allocation to another study.

After families were recruited or allocated to the lab, they were invited to attend an orientation session to provide parents with an overview of the scanning protocol and give them an opportunity to ask questions before providing informed consent. Until the COVID-19 pandemic, and again more recently, orientation sessions were conducted in-person with the aim of acclimating the parent/guardian and infant to the facility and scanner environment. During much of the pandemic, the orientation was conducted online via videoconferencing. Of the 140 families that attended an orientation session, 109 families (77.9%) enrolled and completed at least one scan.

#### Data acquisition

Data were acquired with a 3T Siemens Skyra MRI at Princeton University and a 3T Siemens Prisma MRI at both Yale University locations. The bottom portion of the 20-channel Siemens head coil was used at all locations, with the top portion of the coil removed. Functional images were collected using a whole-brain T2*-weighted gradientecho EPI sequence (Princeton and Yale MRRC: TR = 2000 ms, TE = 28 ms, flip angle = 71°, matrix = 64 × 64, slices = 36, resolution = 3 mm isotropic, interleaved slice acquisition; Yale BIC: identical except TE = 30 ms; slices = 34). Anatomical images were acquired with a T1 PETRA sequence (TR1 = 3.32 ms, TR2 = 2250 ms, TE = 0.07 ms, flip angle = 6°, matrix = 320 × 320, slices = 320, resolution = 0.94 mm isotropic, radial slices = 30,000). Additional methods details and validation of apparatus and parameters have been reported previously (6).

#### Procedure

Scanning sessions were scheduled for a time of day when parents felt the infant would be compliant. Sessions were scheduled for 90 minutes and most commonly began in the morning (e.g., 9:30 or 10:30 AM) or late afternoon (e.g., 3:30 PM). Two members of the research team began preparing for each scan 30 minutes prior to the family’s arrival by assembling the experimental set-up (Figure 1). A third experimenter joined the team for data collection (for additional detail about each experimenter’s role see (6)). Upon arrival, infants, their caregiver, and any toys/comfort items the family brought (e.g., pacifiers, bottles, toys, blankets) underwent extensive metal screening. Infants were shown engaging toys and/or age-appropriate videos to distract them as an experimenter applied three layers of hearing protection: silicon inner-ear putty, over-ear adhesive covers, and earmuffs. The experimenter then laid the infant on the scanner bed and wrapped them in a motion-reducing vacuum pillow with their head in the coil. At least one parent remained in the room for the duration of most scans, except in cases of fMRI contraindication or pregnancy, but they did not enter the scanner bore with the child. At least one experimenter also remained in the scanner room outside of the bore, though often reached in to provide or adjust the infant’s comfort items. Stimuli were projected onto the ceiling of the scanner bore, directly in the infant’s line of sight, via a mirror positioned at the back of the bore (Figure 1B). Infants were monitored using an MRI-compatible video camera (Figure 1C)Princeton and Yale MRRC: MRC 12M-i camera; Yale BIC: MRC high-resolution camera). Video recordings were later used for gaze coding of eye movements. No real-time quality control software was used to quantify infant head motion, but the researcher operating the scanner monitored the video feed to note moments of substantial movement and ensure aliasing was not present in the brain images.

**Fig. 1.**
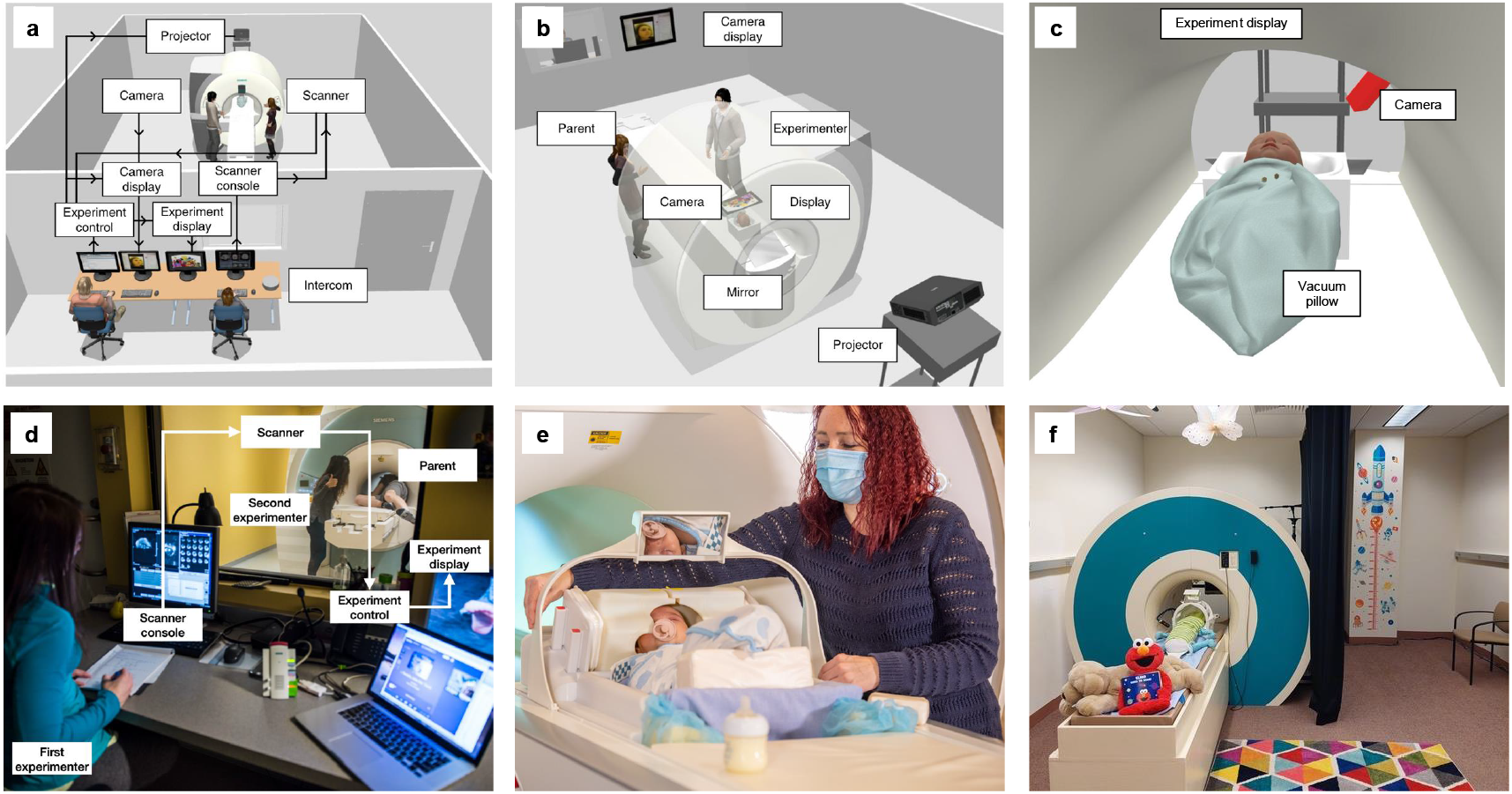
Top panel: Depiction of Yale’s scanning environment including the (A) setup and personnel, (B) projection system for stimulus presentation, and (C) apparatuses used with the infant in the scanner bore. Adapted from (6). Bottom panel: Depiction of MIT’s scanning setup including the (D) infant setup and personnel, (E) custom size-adaptive 32-channel array coil used for infants, and (F) toddler setup before entering the scanner room. Images from Figure 1D and 1E courtesy of Kris Brewer of The Center for Brains, Minds, and Machines.

Stimuli were projected using MATLAB, Psychtoolbox, and in-house ‘experiment menu’ software during functional scans (6). The experiment menu was designed to help researchers navigate the unpredictable nature of infant scanning, allowing for flexible switching from one experiment to another, or from an experiment to a movie used to occupy the infant during anatomical scans. Multiple experiments were prepared in advance for each session, with priority for experiments that the infant did not complete in a prior session. In almost all reported sessions (> 98%), experiments contained visual stimuli only with no auditory stimuli. At least one experiment was attempted before collecting an anatomical image (to be used for alignment); more than one anatomical image was acquired when possible (63 sessions) to ensure that at least one would be of sufficient quality for anatomical alignment during preprocessing. On the rare occasion that usable functional data were collected without a usable anatomical image from the same session (32 sessions), the functional data were excluded. Breaks were taken during or between experiments when the experimenter felt it would improve data quality or when the parent indicated that they wished to pause. These breaks often occurred outside of the scan room so that the infant could feed and/or play.

#### Experimental design

The data for the present analyses were collected during 16 task paradigms that used movie, block, or event-related designs (Table 1). In total, 652 experiments were attempted across the 304 sessions. For present purposes, an experiment was deemed successful if the infant provided enough usable data after exclusions (e.g., head motion, looking behavior) to meet the *a priori* minimum data inclusion criteria of that experiment. Additional experiment-specific counterbalancing factors were ignored as they were highly dependent on experimental considerations and not reflective of the quality of participant data.

**Table 1.**
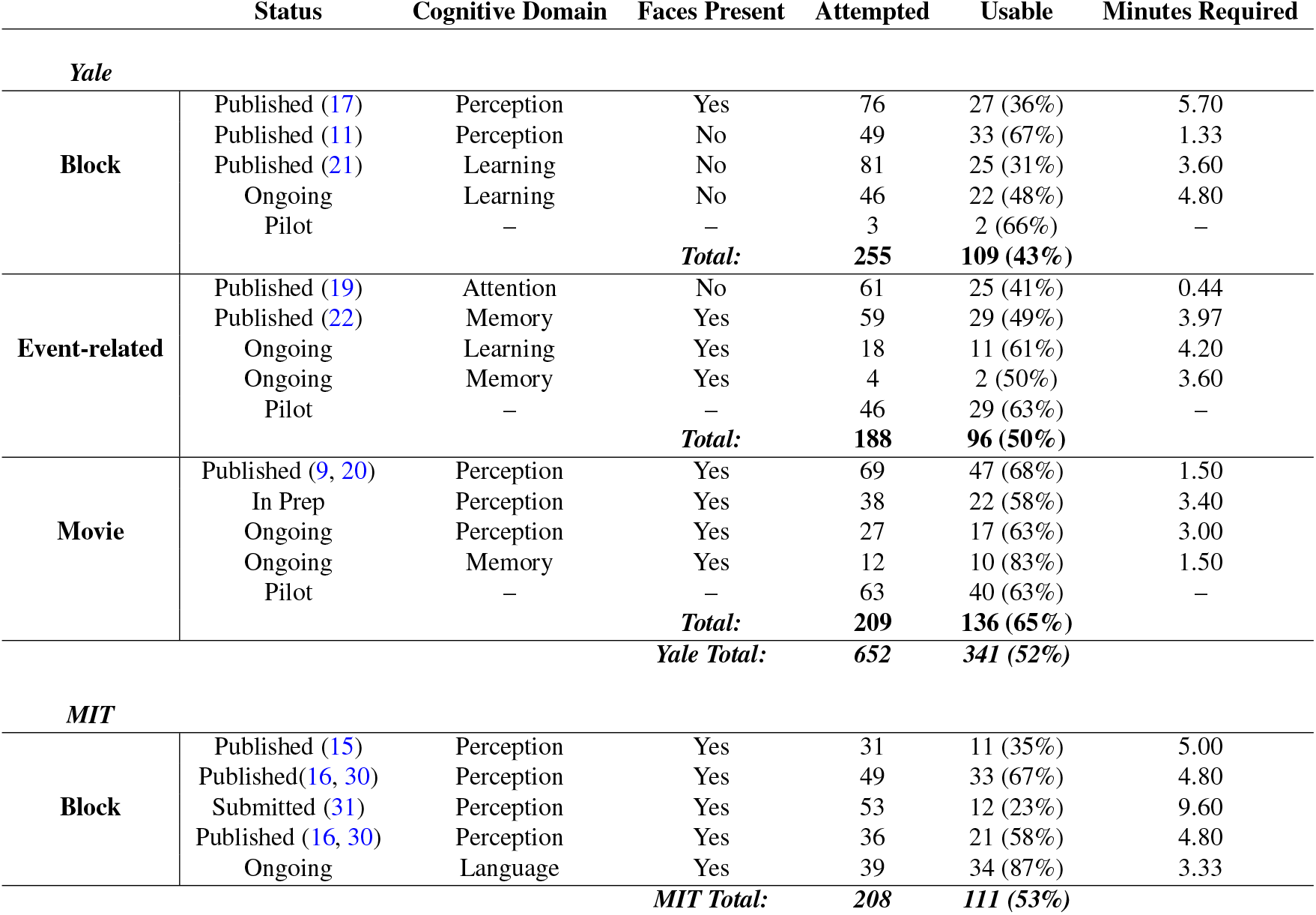
Task datasets collected at Yale from February 2016 to May 2023 and MIT from January 2014 to May 2024.

#### Trait questionnaire

For some infants recruited between February 2016 and February 2021 (*N* = 57), parents completed the Infant Behavior Questionnaire-Revised (IBQ-R). This widely used measure of infant temperament consists of 37 questions assessing three broad dimensions: surgency/extraversion, negative affectivity, and orienting/regulation (32). The questionnaire was typically given to parents at the end of the orientation for them to complete prior to the first scan. Infants ranged in age from 3-11 months old (*M* = 6.15, *SD* = 1.92) at the time the questionnaire was completed. Across the entirety of their participation, these infants attended 175 sessions.

#### State questionnaire

For some infants recruited between October 2021 and May 2023 (*N* = 32), parents completed a Qualtrics survey designed to assess the state of their child at the start of the session. The questionnaire consisted of 16 items assessing the child’s recent mood, sleep schedule, and screen time usage. Families who returned for multiple sessions completed this questionnaire each time (range: 1-5, *M* = 1.91). For present purposes, the following survey items were analyzed: sleep consistency over the last week (3-point scale of “No”, “Somewhat”, or “Yes”); screen exposure, screen interest, and resistance to being laid on back for a diaper change (all using a 5-point scale from “None at all” to “A lot” ); and mood today (7-point scale from “The saddest day” to “The happiest day”).

#### Preprocessing

Data were preprocessed using a publicly released pipeline for awake infant fMRI studies based on FEAT in FSL (6). When multiple experiments were collected in the same functional run, the data were separated to create experiment-specific ‘pseudoruns’ for each task. Three burn-in volumes were removed from the beginning of each run/pseudorun. The reference volume for alignment and motion correction of each run/pseudorun was chosen as the spatial centroid volume. The slices in each volume were realigned with slice-timing correction. Time points with greater than 3 mm of movement from the previous time point (i.e., framewise displacement) were excluded. Missing time points were interpolated to not bias linear detrending but were then ultimately excised from analysis. If more than 50% of the time points in an experimental unit (e.g., event, block, movie, run) were excluded for head motion, the entire unit was excluded.

#### Offline gaze coding

Manual coding of infant gaze was performed offline by two or more coders. The coders indicated whether the infant’s gaze was on- or off-screen (eyes closed or attention diverted away from stimulus), or undetected (obstructed or out of camera’s field of view). Additionally, many experiments required that coders specify where the infant was looking during on-screen periods (e.g., center, left, right). Coders were highly reliable with an average inter-rater reliability of 89% across previously published experiments (range: 79–94%). If the infant was looking for less than 50% of the frames in an experimental unit (e.g., event, block, movie) that entire unit was excluded. In sessions where no eye data were available because of camera malfunction (*N* = 4), the infant was monitored live by an experimenter and no time points were excluded.

### Massachusetts Institute of Technology

#### Participants

The Saxe Lab at the Massachusetts Institute of Technology (MIT) recruited 252 unique infants aged 1–36 months (*M* = 10.85; *SD* = 10.33; 204 female) and attempted a total of 462 scanning sessions. Data collection for the project took place at Athinoula A. Martinos Imaging Center at MIT, an academic research-dedicated center, from January 2014 to May 2024. Infants recruited from the greater Boston, Massachusetts area resided in a racially, ethnically, and socioeconomically diverse community where English is the primary language spoken. A parent and/or legal guardian provided informed consent on behalf of their child. This research was approved by the Institutional Review Board at Massachusetts Institute of Technology.

#### Recruitment

Recruitment methods evolved over time. Early recruitment mainly relied on word of mouth and flyers posted on MIT’s campus, local childcare centers, schools, and community buildings. More recent recruitment was conducted mainly online via social media (Facebook and Instagram) and word of mouth.

#### fMRI data acquisition

Data were acquired with 3T Siemens Trio and 3T Siemens Prisma MRI scanners. For scans conducted from January 2014 to July 2019, a custom 32-channel infant coil for the Trio scanner was used (33). Functional images were collected using a quiet EPI sequence with sinusoidal trajectory (TR = 3s, TE = 43 ms, flip angle = 90°, matrix = 64 × 64, slices = 22, slice thickness = 3 mm; (34)). For scans conducted from late July 2019 to May 2024, a new adjustable 32-channel infant coil was used (28). This coil provided better hearing protection for infants, allowing for a regular EPI sequence with the standard trajectory (TR = 3s, TE = 30 ms, flip angle = 90°, matrix = 80 × 80, slices = 44, slice thickness = 2 mm). Six infants had data collected under a slightly different EPI sequence with more slice coverage (TR = 3s, TE = 30 ms, flip angle = 90°, matrix = 104 × 104, slices = 52, slice thickness = 2 mm). Functional data collected with the new adjustable coil were less distorted than data collected with the initial custom coil. Additional methods, details and validation of apparatus and parameters have been reported previously (16). For participants 18–36 months old, the bottom and top portions of the standard Siemens 32-channel head coil were used with a gradient-echo EPI sequence (TR = 2s, TE = 30.0 ms, flip angle = 90°, matrix = 70 × 70, slices = 46, slice thickness = 3 mm).

#### Procedure

Scanning sessions were scheduled for two hours and session start times were determined based on when parents felt that the children would be most awake and compliant. Two experimenters were present for the session. They arrived and began assembling the scanning environment 30-45 minutes prior to the family’s scheduled arrival. One experimenter greeted the family when they arrived, while the other experimenter finished any setup required. At the start of the visit, participants and their families were introduced to the procedures, consented, screened for metal, and familiarized with the scanner environment. Infants were checked to ensure they were wearing metal-free clothes (i.e., onesies without metal snaps) and adults changed into medical scrubs if they wished to enter the scanning room. Caregivers were encouraged to bring any metal-free toys, pacifiers, bottles, and blankets that they felt might be comforting to their child. Upon entering the scanning room, caregivers distracted the infant as the research team applied the headphone system: over-ear adhesive covers with custom-fit Sensimetrics S15 earbuds, and infant MR-safe earmuffs (Em’s 4 Bubs). Young infants (1–6 months old) were swaddled if possible, or if the caregiver thought it would comfort the child. The experimenters placed the infant supine in the head coil, which was moved into place to fit around the headphones (Figure 1E). Engaging, infant-friendly videos were projected onto a screen at the back of the bore. A mirror positioned directly above the head coil reflected the videos to the infant, while task-irrelevant lullabies were played from the earbuds. Before starting the scan, the experimenter played peek-a-boo with the infant to ensure that the infant could properly see the stimuli at the back of the bore through the mirror. If it seemed the infant might be comforted to have an adult on the bed with them, the caregiver was invited to enter the bore with the infant. If the caregiver did not want to lie in the bore, a researcher lay on the bed instead. During the scan session, a researcher remained in the scanning room for the entire time to communicate with the scan operator, including information about when the infant’s eyes were closed. No real-time quality control software was used to quantify infant head motion. Experimental stimuli were projected to infants using Psychtoolbox.

Procedures for older infants/toddlers (18–36 months) differed from the infant scanning protocol in several ways: A week before the session, parents were sent suggestions of how they could help prepare their child. These instructions included a video to watch together of a child engaging in a rocketship adventure to space, a child-friendly book about the MRI visit, samples of the MRI scanner sounds, and ideas for acclimation to various aspects of the visit (lying down, wearing headphones, etc.). Once participants and their caregivers arrived at a session, they were given time to acclimate to the physical setting and the research team. The acclimation period typically lasted 30–45 minutes. During these older infant sessions, stimuli were projected using Psychopy and task-relevant auditory stimuli were used.

Across all sessions, functional data collection was attempted before an anatomical image was collected. The anatomical scan was started if the participant fell asleep or became fussy during the task; in the latter case, child-friendly videos were played during the anatomical. The experiment could be paused or stopped if the caregiver indicated a need for a break, if infants became too fussy to continue or toddlers asked for a break, and/or when the experimenter felt that a break would be helpful. If the child was removed from the scanner for a break, caregivers were consulted in determining whether it seemed feasible to return to the scanner for a second attempt within the same visit. In many cases, participants and their families returned for multiple sessions.

#### Experimental design

The data for the present analyses were collected in four task paradigms. All used block designs and video stimuli. In three of the task paradigms, infants saw short videos of children’s faces, bodies, objects and natural scenes. Videos lasted 3 s each and were presented in groups of six from the same category, for blocks of 18–20 s. Unrelated music played in the background throughout the experiment. In the fourth task paradigm, participants saw 20-s audiovisual clips of puppets from Sesame Street and heard voices synchronized to the videos. Data collection was considered successful on a given visit if the child provided any usable minutes of data after exclusions (e.g., head motion, eyes closed, awake, etc.). For infants, TRs were excluded because of inattention or sleep based on notes made during the scan session; the researcher inside the scan room communicated the infant’s state to the scan operator. Whether data were usable was defined by the motion thresholds and number of contiguous low-motion measurements required for analysis, which varied between experiments.

#### Preprocessing

Infant data from 2014 to 2022 were preprocessed using a publicly available MATLAB and FSL pipeline (16), while data from 2022-2024 were preprocessed using Python and FSL. For these datasets, consecutive volumes with more than two degrees or 2 mm of motion were cleaved. Only ‘subruns’ with at least 24 consecutive awake, low-motion volumes (those with fewer than than 0.5 degrees or mm of motion between volumes) were retained. Each subrun was then processed individually. One functional image was extracted to be used for registration.

Older infant data (18–36 months) were preprocessed using fMRIPrep 1.2.6 (35). Exclusion thresholds were determined using fMRIPrep Frame Displacement (FD) estimate per run. Volumes within each run that had greater than 1 mm FD or 1.5 standardized DVARS were excluded. If greater than 33% of any run was flagged as motion, the whole run was excluded.

### Present analyses

Data were combined across the two institutions whenever possible, and analyses that could leverage data from both institutions were prioritized. When combining data, there were two key differences between labs which needed to be acknowledged. First, the definition of what constitutes a “session” differed across institutions. In the Yale data, each visit that a family makes to the facility is a new session and there is at least a month between consecutive visits. In the MIT data, all visits a family makes to the scanner within a month are treated as a single session. Although most variables of interest were recorded for each visit (regardless of session definition), two key outcome variables — minutes of data excluded because of motion and minutes of data retained following motion exclusions — could only be calculated on a session-wise basis (i.e., aggregated across multiple visits for some MIT participants). For the main analyses of these outcome variables, the MIT data were limited to sessions in which the participant only visited once in that month, matching Yale’s definition of a session as one visit. In supplementary information, analogous analyses using the MIT definition of a session are reported, retaining all of the Yale sessions (each with one visit) and the single-visit MIT sessions, but now also including the MIT sessions comprised of data from multiple visits within the same month (Supplementary Figure 1).

The second key difference was the motion thresholds that were used during preprocessing (3.0 mm FD and exclusion of runs with more than 50% high motion frames at Yale, 0.5 mm FD and exclusion of subruns with fewer than 24 frames for infants 10 months and younger or 1.0 mm FD and exclusion of runs with more than 33% high motion frames for infants 18 months and older at MIT). These selected motion thresholds are an integral part of each lab’s carefully constructed protocol. It is with these thresholds in mind that each lab designed and refined their experimental practices including their scanning apparatuses, sample sizes, age ranges, task designs, participant scheduling practices, mid-scan decisions about whether to continue or take a break, and more. It would be impossible to disentangle the effects of these other protocol-dependent factors from the impact of the motion threshold itself. Therefore, when reporting on analyses which are impacted by these motion thresholds, the authors include the data as they were collected and preprocessed using each lab’s respective pipelines and thresholds. However, the authors also recognize that the motion thresholds used during preprocessing have important implications for data retention (6). Thus, the authors also replicated analyses after applying MIT’s motion thresholds to Yale’s data. This had the effect of harmonizing the thresholds between Yale and MIT for participants under 10 months (MIT’s infant threshold: 0.5 mm FD and exclusion of subruns with fewer than 24 frames) and over 18 months (MIT’s toddler threshold: 1.0 mm FD and exclusion of runs with more than 33% high motion frames). Notably, MIT did not collect data between 10–18 months, and thus it was unclear which of MIT’s thresholds to apply to Yale’s data from this age range. In the main text, the authors report results with the stricter thresholds applied to these infants (as if younger than 10 months). In the supplement (Supplementary Figure 2), analogous analyses are reported with MIT’s more lenient threshold (as if older than 18 months).

To investigate the factors that predict successful initiation of an awake infant fMRI session, the authors used a logistic regression to model the binary variable of whether an infant entered the scanner and provided a nonzero amount of functional data (yes/no) as a function of their age and institution. The analysis was limited to these predictors because age was the only demographic information collected consistently at Yale when infants did not provide any MRI data. The results of this analysis are not intended to reflect the overall success of a scanning session (which instead depends upon the amount of usable data collected and experimental task criteria), but instead speak to the likelihood of commencing data collection, which posed a significant challenge in many scanning sessions.

For infants who entered the scanner and provided functional data, assigned sex and time of day (hour of scan start time) were also available as predictors. For these infants, multiple regression was used to model the number of minutes of awake functional data collected as a function of institution, age, sex, and hour of scan. A second multiple regression with the same predictors was used to model the number of minutes of functional data retained after preprocessing. Of note, these regression analyses were performed treating each session independently rather than grouping multiple sessions from a given unique infant (when available) in a mixed effects model. This decision was made to avoid assuming that performance should be related across sessions within-subject, given that the additional sessions were often collected after multiple months during which there were rapid developmental changes. Moreover, because only a subset of infants returned, and the interval between sessions was highly variable, there is not adequate data for longitudinal modeling. This approach mirrors the authors’ prior awake infant fMRI publications (11, 17, 19, 21).

Nevertheless, the existence of practice effects in participants who returned on multiple occasions was evaluated. This is a difficult analysis to interpret because of a potential survivor bias. Namely, although successful data collection in one session was not used as a screening criterion for inviting participants back for another session, the families of infants who provided larger quantities of usable data may have been more likely to return for additional scans because of their positive experience, relative to infants who were unable to enter the scanner and/or unable to remain still and complete experiments. As a result, repeat sessions may contain a higher proportion of successful scans, creating a false sense that individual infants yielded more data with repeated testing. To control for this potential bias, a change score was calculated for all pairs of consecutive sessions of an infant by subtracting the number of minutes of data retained after preprocessing in the first session from the number of minutes retained in the second session. In other words, practice effects were evaluated only within individuals who returned. Positive scores indicate improved performance from one session to the next. A multiple regression model was used to test for effects of session number and infant age on the change scores.

In addition to predicting overall minutes of usable data retained (which relied on different motion thresholds from each institution), multiple regression was used to predict two key sources of data loss: excess head motion (reported by both institutions) and visual inattention (looking away from the stimulus or closing eyes; reported by Yale). The proportion of data excluded for excess motion and visual inattention were modeled as a function of age, sex, hour of scan, and institution (when appropriate).

For the state questionnaire collected at Yale, multiple regression was used to test for associations between parent-report items and usable minutes of functional data on the day that the questionnaire was completed. For the trait questionnaire, Pearson correlation was used to test how minutes of usable functional data for each session related to their score on three temperament summary variables: surgency/extraversion, negative affectivity, and orienting/regulation (32).

Beyond these factors centered on the infant, the Yale data permitted consideration of how experimental practices impacted scanning success. The efficacy of attempting to collect multiple experiments during each scanning session was quantified based on the number of experiments that yielded sufficient usable fMRI data to be considered complete. The success rates of different experimental designs and stimulus contents were compared with a chi-square test.

## Results

### Awake functional data collected

#### Likelihood of collecting any functional data

Data from 683 sessions were available to assess the binary likelihood of collecting functional data during a session (304 Yale, 379 MIT; Figure 2, Step 1). In most cases (*N* = 498, 72.9%), the infant successfully entered the scanner and provided a nonzero amount of functional data (age: *M* = 9.59 months, *SD* = 7.95). In the other sessions (*N* = 185, 27.1%), no functional data were acquired (age: *M* = 17.14 months, *SD* = 9.64). These failed scans occurred for a variety of reasons, including: non-compliance with hearing protection, refusal to separate from caretaker, and unwillingness to remain lying down once in the scanner bore. A multiple logistic regression model predicting whether any functional data were collected (yes/no) as a function of institution and infant age (Figure 3A-C) provided a significant overall fit (*χ*^2^(3) = 117.62, *p* < .001). There was a greater likelihood of collecting functional data from younger infants (*β* = -0.08, *p* < .001; odds ratio [OR] = .92, 95% confidence interval [CI] = [.90, .94]) and from infants scanned at Yale (*β* = -1.06, *p* < .001; OR = .35, 95% CI = [.24, .52]).

**Fig. 2.**
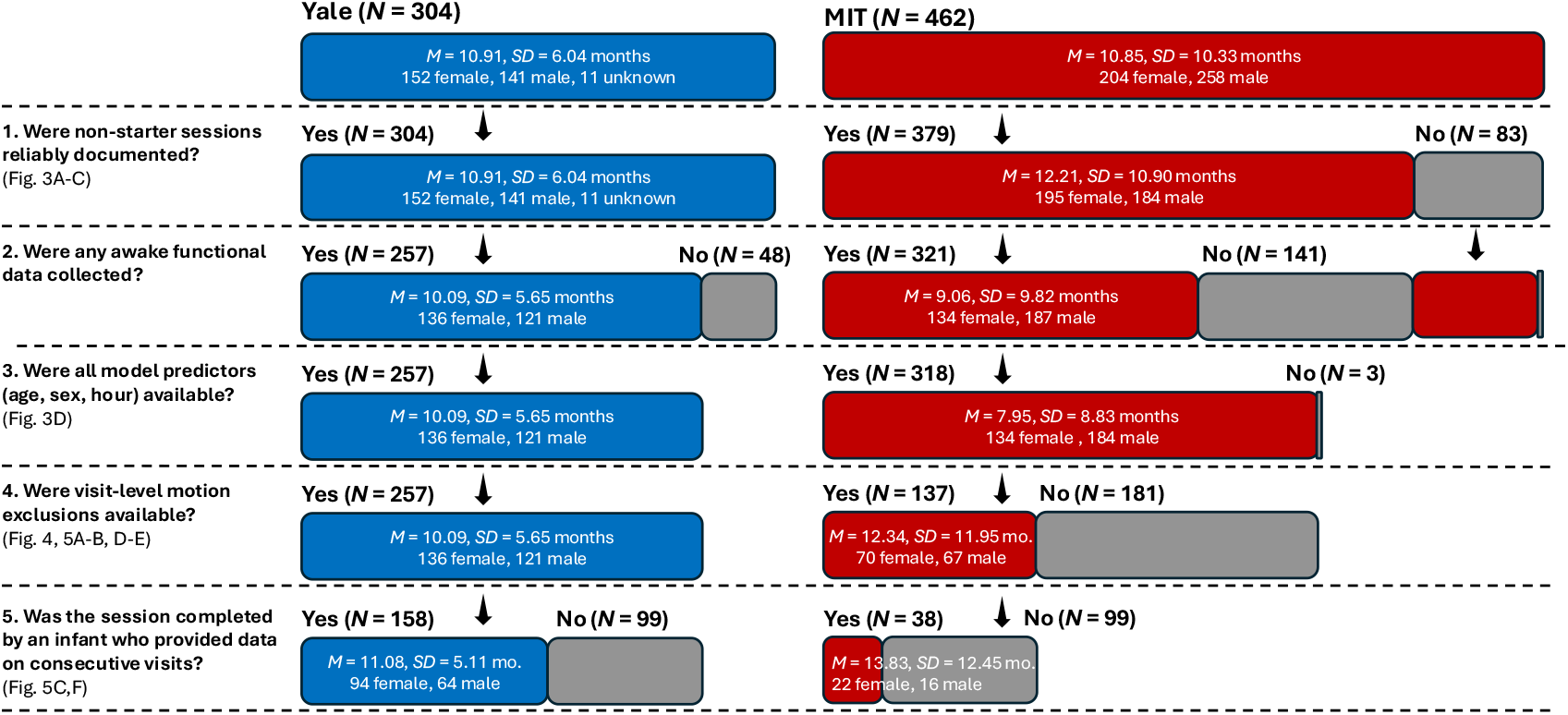
Overview of participant inclusion criteria by institution. For each analysis step, the pool available from the previous step is divided into those who are included (Yes) or excluded (No). Available demographic information is depicted for included participants. Bar size is proportional to the number of participants represented.

**Fig. 3.**
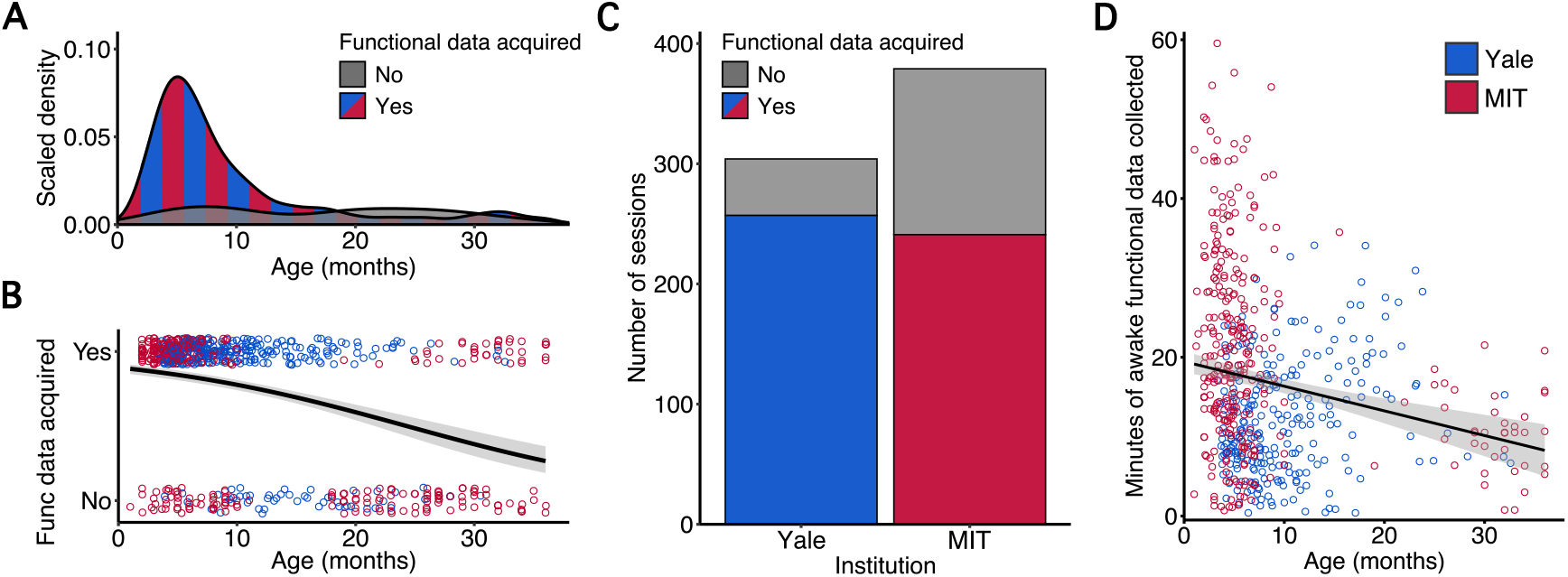
Factors predicting functional data collection. (A) Density plot separated by infant willingness to enter the scanner, as a function of age in months. The colored curve shows the density across ages of infants who did enter the scanner and provide some amount of functional data combining across both institutions; the gray curve shows the density of those who did not. (B) Logistic regression predicting whether any functional data were acquired based on infant age in months. Each circle is one fMRI session colored by the institution where it was conducted, the black line is the best logistic fit, and the shading is the 95% confidence interval (CI) of the fit. (C) Number of sessions with and without some amount of functional data collected, separated by institution. The colored portion of the bars represents the number of sessions where the infant did enter the scanner and provide some amount of functional data, the gray portion represents the sessions where the infant did not. (D) Scatter plot minutes of awake functional data collected by infant age in months. Each circle is one session, the black line is the best linear fit, and the shading is the 95% CI of the fit.

#### Minutes of awake functional data collected

Data from 575 sessions where the infant entered the scanner and provided some amount of functional data were available to assess factors impacting the *amount* of data collected (257 Yale, 318 MIT; Figure 2, Step 2–3). In these sessions, an average of 16.69 total minutes (Range = [0.40 – 59.55], *Mdn* = 14.67, *IQR* = 14.40) of functional data were collected. A multiple regression model predicting the number of minutes of functional data collected during a session as a function of institution, infant age, assigned sex, and hour of scan provided a significant overall fit (*F*(4, 570) = 26.00, *p* < .001, *R*^2^ = .15, *R*^2^adj = .15); institution (*β* = 7.42, *p* < .001; 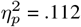) and infant age (*β* = -.25, *p* < .001; 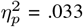) were significant predictors. Namely, younger infants and sessions conducted at MIT were associated with more minutes of data collected (Figure 3D). These same predictors were significant when the model was tested after applying a square-root transformation to the dependent variable to account for the skewed distribution. Post-hoc analyses were conducted to verify the linearity of the age effect. Bayesian information criterion (BIC) for the model with a linear age term (4,335.93) was slightly lower than for a model with linear and quadratic age terms (4,336.73), suggesting that the linear model better balanced fit and complexity.

### Data excluded during preprocessing

#### Proportion of data excluded because of head motion

Data from the 394 sessions during which infants provided some amount of functional data and head motion was calculated on a visit-wise basis were used to investigate factors impacting data exclusion (257 Yale, 137 MIT; Figure 2, Step 4). The most common cause for exclusion was excessive head motion, leading to an average loss of 5.74 minutes (39.4%) of functional data per session. This rate depended heavily on an experimenter-defined, *a priori* framewise displacement threshold that differed by institution (3 mm at Yale, 0.5 or 1.0 mm MIT), as shown previously (6). Therefore, the authors first examined how various factors affected the amount of data loss, with each lab using the threshold they have each adopted in their respective publications. A multiple regression model of the proportion of data excluded because of head motion within a session as a function of institution, infant age, assigned sex, and hour of scan provided a significant overall fit (*F*(4, 389) = 65.84, *p* < .001, *R*^2^ = .40, *R*^2^adj = .40). Specifically, infant age was a significant predictor (*β* = -0.01, *p* < .001; 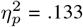), with more minutes excluded for younger infants (Figure 4A, Left Panel), and institution was a significant predictor (*β* = 0.42, *p* < .001; 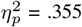), with more minutes excluded at MIT (Figure 4B, Left Panel). The authors also conducted this analysis after applying MIT’s motion thresholds to Yale’s data. This model provided a significant overall fit (*F*(4, 389) = 21.26, *p* < .001, *R*^2^ = .18, *R*^2^adj = .17). In addition to age (*β* = -0.02, *p* < .001; 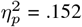; Figure 4A, Right) and institution (*β* = 0.08, *p* = .016; 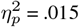; Figure 4B, Right), assigned sex was also a significant predictor (*β* = -0.08, *p* = .010; 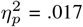). Thus, although the stricter thresholds closed the gap in motion exclusion between labs (as expected), the effects of age and institution persisted in the harmonized data; interestingly, a sex difference emerged that was not reliable before (more exclusion in males).

**Fig. 4.**
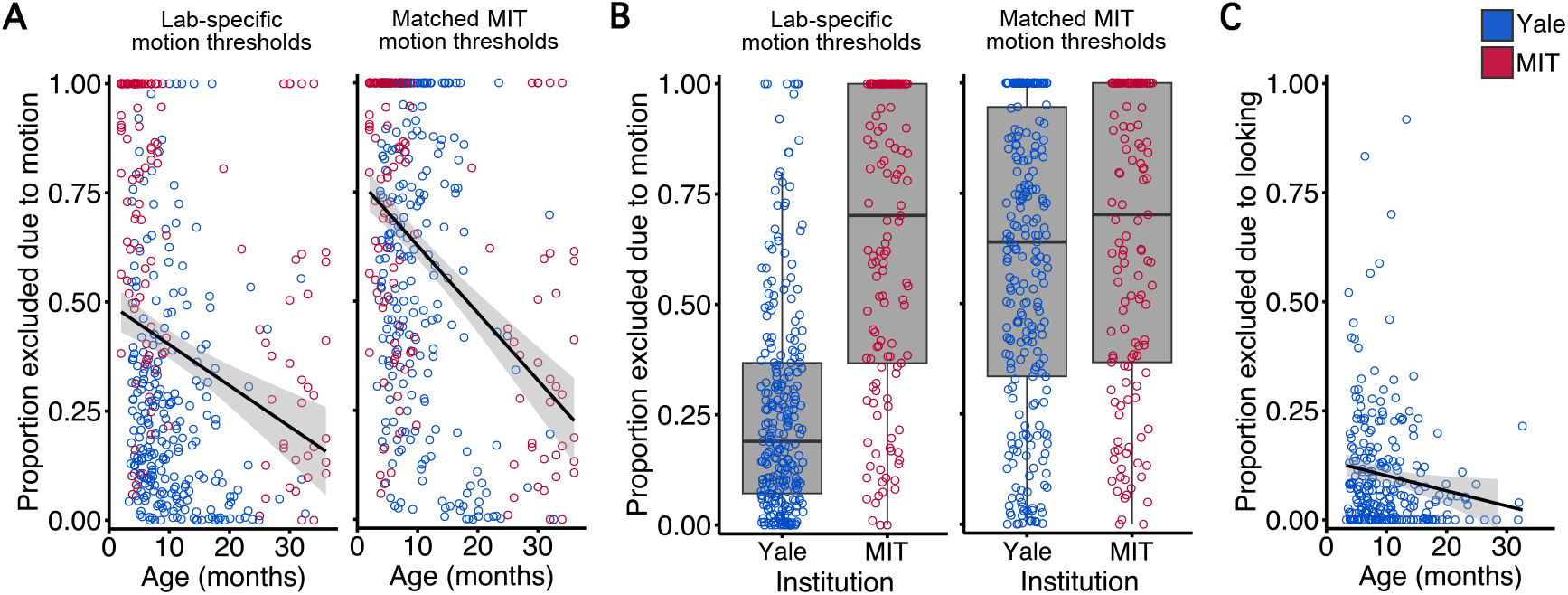
Factors predicting functional data exclusion. (A) Scatter plot of proportion of functional data excluded because of head motion by infant age in months. Each circle is one session colored by institution, the black line is the best linear fit, and the shading is the 95% CI of the fit. Left: Motion exclusions calculated using Yale’s (3 mm) and MIT’s (0.5 mm for 1–10 months, 1 mm for 18–36 months) respective thresholds. Right: Motion exclusions for both institutions calculated using MIT’s motion thresholds (0.5 mm for 1–18 months, 1 mm for 18–36 months). (B) Proportion of functional data excluded because of head motion by institution. Each circle is one session and the box plot depicts the distribution across four quartiles. Left: Motion exclusions calculated using the respective thresholds of Yale and MIT. Right: Motion exclusions for both institutions calculated using MIT’s motion thresholds. (C) Scatter plot of proportion of functional data excluded because of visual inattention (i.e., not looking to stimulus display) by age in months (only available for Yale). Each circle is one session, the black line is the best linear fit, and the shading is the 95% CI of the fit.

#### Proportion of data excluded because of visual inattention

Among the functional data retained after head-motion exclusions, an additional 1.04 minutes (8.4%) per session were excluded on average from the Yale sessions (*N* = 257) because of visual inattention (i.e., gaze directed away from the stimulus or eyes closed). A multiple regression model of proportion of data excluded because of visual inattention as a function of infant age, assigned sex, and hour of scan also provided a significant overall fit (*F*(3, 253) = 2.69, *p* = .047, *R*^2^ = .03, *R*^2^adj = .02). The effect of infant age was significant (*β* = -0.003, *p* = .029;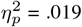), with a greater proportion excluded for younger infants (Figure 4C).

### Data retained after preprocessing

#### Minutes of usable functional data retained

Data from 394 sessions were available to assess factors impacting the amount of usable data retained after motion exclusions (257 Yale, 137 MIT; Figure 2, Step 4). In these sessions, an average of 14.58 minutes of functional data were collected, of which 8.82 (60.4%) minutes were retained (Range = [0 – 36.60], *Mdn* = 6.92, *IQR* = 11.22). A multiple regression model predicting the number of minutes of usable functional data retained from a session as a function of institution, infant age, assigned sex, and hour of scan provided a significant overall fit (*F*(4, 389) = 7.66, *p* < .001, *R*^2^ = .07, *R*^2^adj = .06); infant age (*β* = 0.10, *p* = .038; 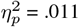), assigned sex (*β* = 1.71, *p* = .026; 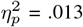), and institution (*β* = -3.57, *p* < .001; 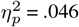) were significant predictors. This model with a linear age term had a higher BIC (2,740.57) than a model with linear and quadratic age terms (2,731.85). The fit of this linear+quadratic age model was unsurprisingly also significant (*F*(5, 388) = 9.29, *p* < .001, *R*^2^ = .11, *R*^2^adj = .10), with the quadratic term for infant age (*β* = -33.84, *p* < .001; 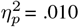) and assigned sex (*β* = 1.50, *p* = .047; 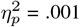) as significant predictors. Namely, infants provided more usable data between 12–24 months old than 0–12 or 24–36 months old (Figure 5A), and female infants provided more usable data than male infants (Figure 5B). These same predictors were significant when the model was tested after applying a cube-root transformation to the dependent variable and removing values at floor (0 minutes of usable data) to account for the non-normal distribution. A model fit with an additional term for an interaction between age and assigned sex (BIC = 2,737.05) indicated there was no such interaction (*p* = .381).

**Fig. 5.**
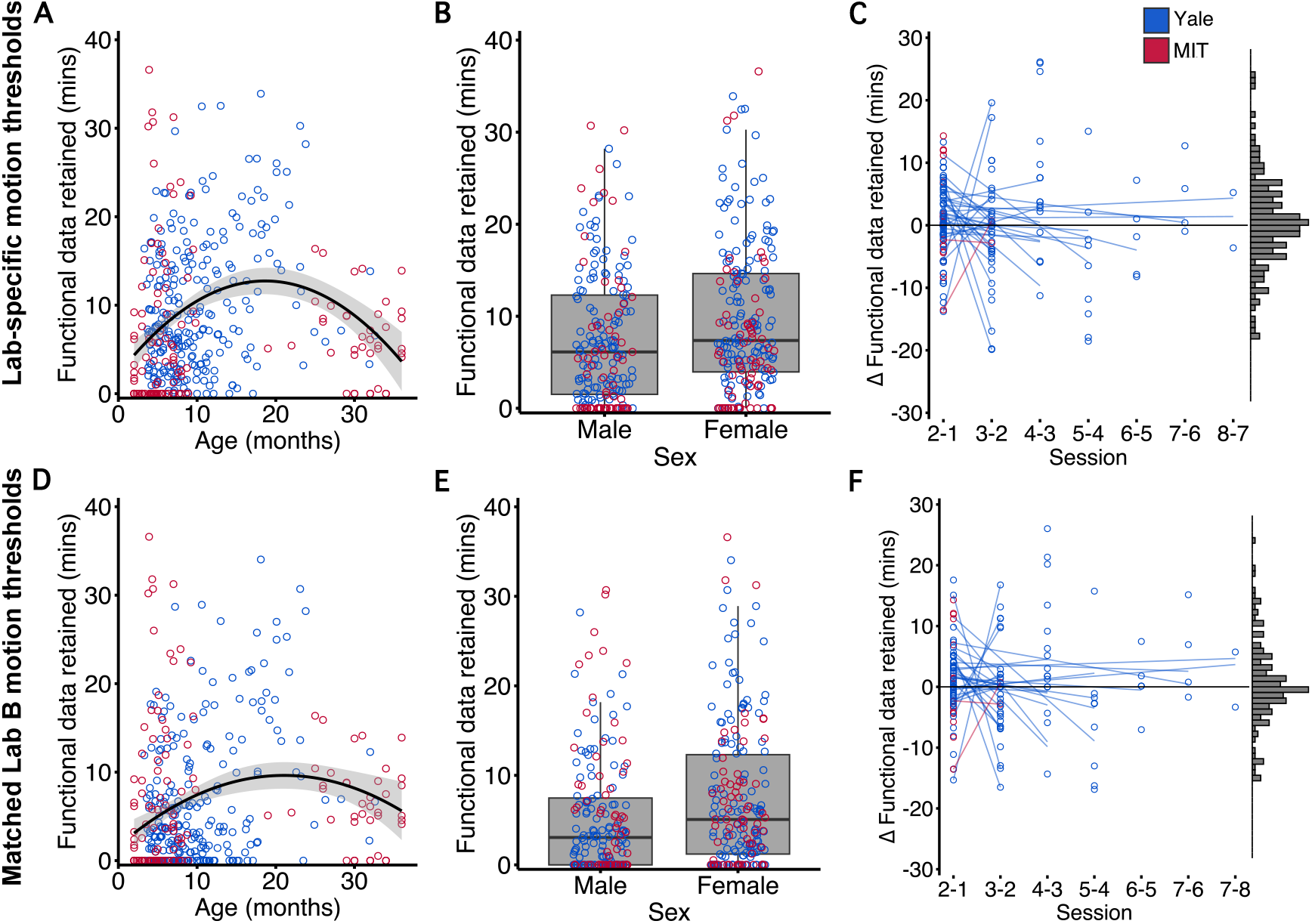
Factors predicting functional data retention. Top row: Minutes of data retained calculated based on each lab’s respective thresholds (Yale: 3.0 mm; MIT: 0.5 or 1.0 mm). (A) Scatter plot of minutes of functional data retained after motion exclusions by infant age in months. Each circle is one session colored by institution, the black line is the best linear+quadratic fit, and the shading is the 95% CI of the fit. (B) Minutes of functional data retained by sex assigned at birth. Each circle is one session and the box plot is the distribution across four quartiles. (C) For repeat participants, the change in minutes of functional data retained for the current session minus the previous session. The solid lines indicate the best linear fit for each participant; longer lines indicate participants who returned for more sessions. The marginal histogram shows the overall distribution of the change scores; the centering at zero indicates the lack of a practice effect. Bottom row: Minutes of data retained based on harmonizing to MIT’s thresholds (0.5 mm for 1–18 months, 1.0 mm for 18–36 months). (D) Scatter plot of minutes of functional data retained after motion exclusions by infant age in months. The black line is the best linear+quadratic fit, and the shading is the 95% CI of the fit. (E) Minutes of functional data retained by sex assigned at birth. Each circle is one session and the box plot is the distribution across four quartiles. (F) The change in minutes of functional data retained for the current session minus the previous session. The solid lines indicate the best linear fit for each participant. The marginal histogram shows the overall distribution of the change scores. See also Supplementary Fig. 2.

The authors also conducted this analysis after applying MIT’s motion thresholds to Yale’s data. In this case, an average of 6.49 (44.5%) minutes of functional data were retained (Range = [0 – 36.60], *Mdn* = 3.83, *IQR* = 9.71). The same multiple regression model with linear and quadratic age terms provided a significant overall fit (*F*(5, 388) = 6.46, *p* < .001, *R*^2^ = .08, *R*^2^adj = .06). In addition to the quadratic term for age (*β* = -33.46, *p* < .001; 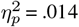; Figure 5D) and assigned sex (*β* = 1.80, *p* = .016; 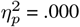; Figure 5E), the linear age term was also a significant predictor (*β* = 18.87, *p* = .012; 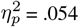); as before, institution was not a significant predictor.

#### Change in amount of usable data across repeat sessions

Many infants from each lab attended multiple scanning sessions. At Yale, of the 115 unique infants that have participated, 75 (65.2%) elected to return and complete two or more scanning sessions. At MIT, of the 252 unique infants that have participated, 112 (44.4%) completed two or more scanning sessions. Data from 196 sessions (using the Yale definition of a session; 158 Yale, 38 MIT) in which infants returned were available for the present analysis of whether there was a practice effect in the amount of usable data as a function of session number (Figure 2, Step 5). To test whether prior scanning experience impacted the amount of usable data, change scores were computed for each pair of sessions completed by a returning infant as the number of minutes of data retained in the current session minus minutes retained in the previous session. The distribution of pairwise change scores (Figure 5C, right; *M* = 0.43, *SD* = 8.16) did not significantly differ from zero (*t*(151) = 0.65, *p* = .514), inconsistent with an overall practice effect. In addition, a multiple regression model was used to predict the change scores as a function of session number and infant age. There was no significant main effect of session number (*β* = 0.27, *p* = .612) or infant age (*β* = -0.14, *p* = .215), suggesting that a practice effect did not emerge with more repetition or in older infants (Figure 5C, left). These results were similarly null when assessed after MIT’s thresholds were applied to Yale’s data (Figure 5F).

### Experiment and participant factors impacting data retention

#### Benefit of attempting multiple experiments

Given the massive variability in usable functional data collected per session, the authors tested the efficacy of attempting to collect multiple experiments per session at Yale. Across all 304 Yale sessions, 652 experiments were attempted and 341 (52.3%) of these were usable after preprocessing. This corresponds to an average of 2.14 attempted and 1.12 usable experiments per session. The 341 usable experiments came from 178 of the 304 sessions. As such, at least one usable experiment was obtained in 59% of sessions (Figure 6A). Despite 41% of sessions yielding no usable experiments, the 1.12 usable experiments on average across all sessions occurred because a majority of the successful sessions yielded more than one usable experiment (104 of 178). In other words, 47.8% of usable experiments were not the first usable experiment collected during the session and would not have been completed if data collection had stopped after a single experiment.

**Fig. 6.**
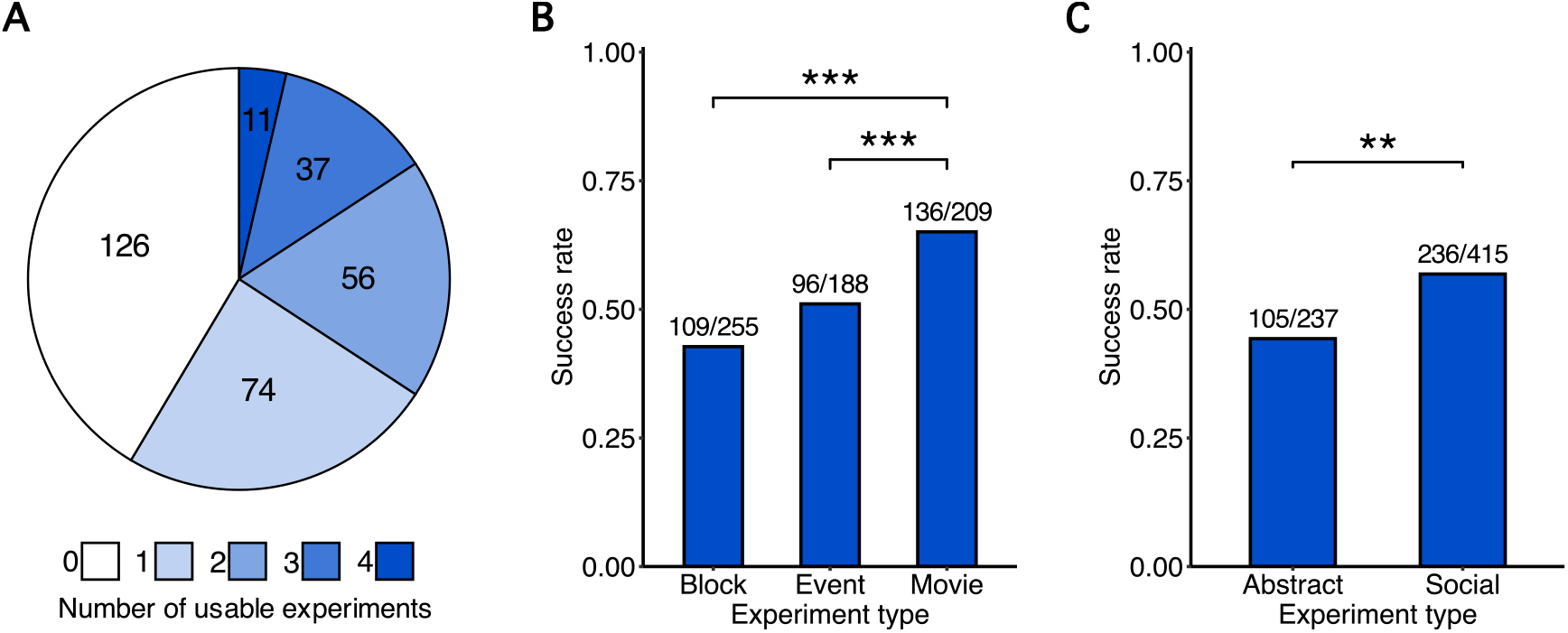
Experimental factors relating to scan success. (A) Number of usable fMRI experiments acquired during each Yale session. (B) Success rate (number of usable experiments divided by number of attempted experiments) as a function of experimental task design. *** = *p* < .001 (C) Success rate (number of usable experiments divided by number of attempted experiments) as a function of stimulus content. *** = *p* < .01

#### Experimental design of tasks

The functional data collected at Yale could be categorized into three types of fMRI task designs: movie, block, and event-related (Figure 6B). A chi-square test of independence showed that success rate — the number of usable experiments divided by number of attempted experiments — differed by task design (*χ*^2^(2) = 23.11, *p* < .001). Post-hoc pairwise comparisons using a Bonferroni correction indicated that movie designs had a higher success rate (65.1%) than both block designs (42.7%; *p*adj < .001) and event-related designs (51.1%; *p*adj = .002); the difference between block and event-related designs was not significant (*p*adj = .304). A post-hoc analysis to test whether the effect of experiment type varied across infant age compared a logistic regression with age and experiment type as predictors to one that also included an interaction term. The interaction term did not improve model fit (*χ*^2^(1) = 2.76, *p* = .251), suggesting that the effect of experiment type did not differ significantly across ages.

Tasks could also be categorized based on the content of their stimuli. Many different stimuli were used across experiments, but one common dimension is whether they featured social stimuli (e.g., realistic or cartoon faces of any kind) or abstract stimuli (e.g., colorful shapes and textures). A Chisquare test with Yates’ continuity correction showed that success rate differed based on this dimension of stimulus content (*χ*^2^(1) = 9.05, *p* = .003). Namely, tasks featuring social stimuli had a higher success rate (56.9%) than those featuring abstract stimuli (44.3%; Figure 6C). A post-hoc analysis to test whether the effect of stimulus content varied across infant age compared a logistic regression with age and stimulus content as predictors to one that also included an interaction term.

The interaction term did not improve model fit (*χ*^2^(1) = 0.30, *p* = .582), suggesting that the effect of stimulus content did not differ significantly across ages.

#### Trait and state characteristics of participants

Trait and state questionnaires were administered to a subset of parents with the goal of uncovering pre-scan factors about the infant that might be useful for predicting successful data collection. For the IBQ-R infant trait questionnaire, there was a significant relationship between negative affectivity and minutes of usable data after Bonferroni correction (*r*(173) = -.25, *p*adj = .002), such that infants with greater reported affectivity provided fewer minutes of usable data. There was no significant relationship between minutes of usable data and surgency/extraversion (*r*(173) = -.04, *p*adj = 1.00) or orienting/regulation (*r*(173) = .07, *p*adj = 1.00). A post-hoc analysis found no significant difference in negative affectivity between males (*M* = 3.38) and females (*M* = 3.50; *t*(52) = -0.46, *p* = .644).

For the infant state questionnaire, a multiple regression model predicting minutes of usable data from sleep consistency, screen exposure, screen interest, mood, and resistance to being laid on back did not fit reliably overall (*F*(5, 55) = 1.58, *p* = .180, *R*^2^ = .13, *R*^2^adj = .05). Despite this, screen interest was a reliable predictor (*β* = -2.26, *p* = .021), with infants reported to have less interest in screens at home providing more usable functional data.

## Discussion

The data that the authors’ labs have collected over the past decade demonstrate the feasibility of awake infant fMRI and provide information about what other researchers might expect when using this method. Across both labs, almost 9 minutes of usable data were collected per session on average. At Yale, the authors collected one or more usable experiments in 58% of sessions. Critically, this success rate is surprisingly similar to reported rates for fNIRS (46–66%) — a technique often considered more infant-friendly (36, 37). Furthermore, Yale averaged 1.1 usable experiments per session, suggesting that attempting to collect multiple experiments from cooperative infants can offset the time and resources spent on unsuccessful sessions.

The authors also demonstrate that age, sex, data collection practices, and experimental designs all impact the success of awake infant fMRI scans. These findings speak to some steps researchers might take to improve the retention of awake infant fMRI data.

### Participant characteristics

The data presented here suggest that future studies should closely consider age when designing experiments. Younger infants are more likely to enter the scanner and provide some amount of functional data; this may make it easier to collect the larger samples needed for between-subject and individual-differences designs. In contrast, despite higher attrition, older infants up to two years produce a greater amount of usable functional data per session after exclusions, which may be important for within-subject designs. Of course, which age to target is principally governed by the developmental question of interest, but knowledge of these constraints may be helpful in designing more effective experiments. It is worth noting in this context that infants under one year of age tend to be oversampled in infant neuroimaging studies across modalities (38).

Assigned sex was also a predictive demographic characteristic, with female infants providing more usable functional data than their male counterparts on average. Of course, this does not mean that studies should not strive for gender balance in their sample. However, it is interesting to consider what might be driving this effect. One potential explanation could be trait-level differences across the sexes, which have been suggested to emerge early (39). However, there was no sex difference in negative affectivity (40) — the only trait related to scanning success in the overall sample. There was also no interaction between sex and age, inconsistent with an explanation based on experiential differences or acculturation. A promising direction could be to consider sex differences in the physiological reactions and stress responses of infants (41). Alternatively, this effect may not reflect an inherent difference between female and male infants themselves, but rather a difference in how researchers and/or caregivers interact with female versus male participants.

However, the parent-report behavioral measures from the infant state questionnaire administered at Yale did not significantly predict the amount of usable functional data overall. The null results for most state and trait measures speak to the unpredictable nature of scanning infants, but should also be interpreted with caution given the limited data available for infants who did not enter the scanner and provide functional data. Moreover, factors not thoroughly assessed in our questionnaires such as teething, hunger, stranger anxiety, and illness all have the potential to unduly influence a scan. For this reason, we encourage all interested families to enroll and attend sessions, even if they feel their baby is unlikely to perform well on a given day. Anecdotally, many sessions have been successful when it seemed unlikely at the start, and unsuccessful even when the circumstances seemed ideal. The fact that an infant’s performance in one session did not predict their performance in subsequent sessions, suggests that it is not the case that one subset of well-behaved infants provides all of our data and a second set of non-cooperative infants are never included. Furthermore, the studies conducted by both labs utilize within-subject designs which are less sensitive to selection bias or cohort effects than between-subject or individual differences designs. That said, there may have been some (self-)selection bias in which families chose to enroll in the first place; indeed, many of the parents whose infants participated were highly educated and often affiliated with our own, or nearby, universities.

### Protocol considerations

There were differences in the amount of data collected and excluded across the two institutions. The goal of this work was not to determine which lab’s approach was superior — indeed, there are lessons and strengths to be taken from each — but these differences emphasize the importance of exploring the impact of lab-specific practices. Most notably, MIT collected significantly more data per session on average (over 20 minutes, compared to 12 minutes at Yale). This difference was partly attributable to the fact that at least one anatomical image was collected in every Yale session, meaning that there were at least three additional minutes of engagement that could have been used to collect more functional data; MIT collected an anatomical image if the infant fell asleep or, in some cases, if the infant seemed to prefer a non-experimental video. Additionally, Yale scanned many more 10–20 month-old infants than MIT, who focused primarily on infants under 10 months. The results from the combined dataset suggest that this may decrease the total amount of functional data collected on average by Yale (and increase the relative amount of those data that were usable).

Moreover, certain details of the MIT protocol may have facilitated compliance to allow for longer periods of data collection. For example, infants scanned at MIT may have been soothed by music played over the headphones and/or the presence of the caregiver/researcher accompanying them in the scanner bore. Additionally, the custom head coil used by MIT places the infant in a half-seated position which may have also increased comfort and led to greater compliance once the session started.

On the other hand, Yale’s use of only the bottom portion of a 20-channel head coil in an open-face configuration was less confining, and the ceiling projection may have engaged the infant’s attention more rapidly, reducing the proportion of sessions in which no functional data were collected. This approach may also be more easily adopted by researchers new to awake infant fMRI, as it relies on stock equipment commonly used with adults. Additionally, the authors have previously shown that this approach produces a comparable signal-to-fluctuation-noise ratio (SFNR) to that obtained when scanning adults with both portions of the coil, particularly in cases of low head motion (6).

Overall, despite the protocol differences between Yale and MIT, the authors also realized that they had developed some similar strategies over time. For example, both groups found that parent comfort with the scanning process was critical for a successful scan. We both found it beneficial to schedule sessions with ample time for acclimation to the environment and breaks during scanning to ensure the process did not feel rushed for participants. Additionally, though each lab used different scanning sequences and forms of hearing protection, we often found that infants were calmer while the scanner was running, so it was best to minimize time spent in the bore with the scanner idle. Though only anecdotal, we have presumed this is due the rhythmic vibrations they felt and/or muffled sounds they heard.

Relatedly, one aspect of MIT’s protocol that Yale is currently working to adopt is this presentation of auditory stimuli to infants while in the scanner. This not only broadens the scope of possible experiments to include auditory/multisensory perception and various aspects of language, but may also be calming to the infant and increase data quality/quantity. Conversely, one strength unique to Yale’s protocol is the video camera positioned inside the scanner bore which allows for tracking of the infant’s eye movements during a scan. Eye movements have been a vital tool for researchers of infant cognition, and monitoring this looking time behavior in the scanner enables investigation into how it relates to brain activity.

Additionally, even though the amount of data retained after motion exclusions was highly variable across sessions, attempting to collect multiple experiments in each session at Yale helped mitigate this variability. In fact, more than one usable dataset was obtained per session on average, despite the fact that almost half of sessions resulted in no usable experiments. The practice of preparing multiple experiments in advance of each session not only increased the amount of data collected from compliant infants, but also allowed for flexibility in adapting to infant interest and task compliance. For example, it was not unusual for an infant to fuss out of one task but then to re-engage with, and complete, a second or third task.

Finally, there are a number of challenges for which both labs are still working to find a solution. For example, as is the case in many types of developmental research, too many sessions fail before any data are collected. In our experience, this is commonly due to fussiness and/or non-compliance with hearing protection, which can be especially difficult with infants who are teething or congested. Over-the-ear noise canceling headphones that reliably and automatically cancel scanner noise may facilitate the application of hearing protection. Moreover, the head motion encountered when scanning infants limits the feasibility of tasks and analyses that require long stretches of usable data (Supplementary Figure 3). This challenge is currently addressed by scanning infants while they are asleep, but this state is not suitable for many questions about cognition.

### Experimental design

Across different tasks at Yale, movie designs were more successful than block and event-related designs on average; MIT only used movie designs. This advantage may partly come from the fact that movie designs tend to require fewer minutes of data to be usable overall, given the continuous stimulation. These findings align with prior demonstrations that movies are a powerful tool for collecting high-quality fMRI data in young children (42). The value of utilizing naturalistic stimuli during fMRI studies is also well established in adults (43). For research questions that require a more traditional task, block and event-related designs yielded similar amounts of usable data; this contrasts with the observation of higher attrition rates for event-related than block designs in fNIRS (36). One potential limitation when interpreting the lack of a difference here is that Yale ran more block designs in earlier years and more event-related designs in later years, thus confounding this comparison with experimenter experience.

The type of stimuli shown within a task was also related to the success rate of experiments run at Yale. Specifically, tasks featuring social stimuli (i.e., faces) had a significantly higher success rate than those featuring abstract stimuli (i.e., shapes, textures); notably, all of the experiments run at MIT involved social stimuli. These results align with prior evidence demonstrating that infants preferentially orient to faces over non-face objects (44). This preference, which increases with age, facilitates social attention (45). Taken together, these findings suggest that social stimuli can be leveraged in future awake infant fMRI studies to facilitate infant orienting and attention to the stimulus display and maximize experimental success rate.

Choices about experimental design and stimuli should also be informed by other aspects of the protocol that impact data quality and quantity. Testing infants in an awake state makes a broad range of tasks and behaviors possible but results in more head motion than when they are asleep. This is often an acceptable compromise because carefully balanced within-subject manipulations make task fMRI less susceptible to motion confounds than resting-state fMRI. Decisions about how to handle head motion during preprocessing further interact with experimental choices. For example, movie designs may be more compatible with a stricter motion threshold than block or event-related designs because their higher success rate offsets the increase in data exclusion. Additionally, the data exclusion associated with a stricter motion threshold can be addressed by attempting the task repeatedly across multiple visits scheduled close in time (when the infant is roughly the same age). MIT has found that this approach works well, especially for cognitive domains such as visual perception and language in which tasks can be repeated easily. Other cognitive domains, such as learning or memory, may not be as amenable to repeated testing (e.g., because the amount of exposure is often part of the phenomenon of interest or there can be interference across sessions), and some questions require block or event-related designs that have lower success rates. In these cases, Yale has found that a more lenient threshold that maximizes the data retained from a single visit can be effective. The resulting reduction in data quality requires simple and robust experimental contrasts and rigorous statistical analyses to find the signal in noise. Imaging older infants, for whom a smaller proportion of data is excluded due to motion, may also help to combat potential concerns of data loss at either threshold.

### Limitations

The scope and generalizability of the reported analyses is limited by the variables that were documented and the research practices that were employed by each institution. There are many factors not considered here that could play a vital role in the scanning process. For example, hearing protection is difficult when a child is teething and some children seem calmer and more amenable to scanning when their guardian is not visible. Although such occurrences are noted, the potential influence of these factors is at best anecdotal. Additionally, while both labs ensure that parents are fully informed and oriented prior to scanning, a relatively small proportion of the sessions reported involve pre-scan accommodation/preparation for the subject (only for 18–36 month-olds at MIT). This can be helpful at these older ages, but it is also possible that taking analogous steps may be beneficial for younger infants, potentially by easing the parent’s anxiety as much as the child’s (46). Indeed, the behavioral questionnaires reported here focused entirely on the infant, yet it is possible that parent characteristics may play a strong role. Protocols for infant/toddler neuroimaging vary dramatically across labs (29), and it will be important to assess these divergent approaches and glean insights from these groups — including those using other infant imaging modalities — for establishing best practices.

## Conclusion

Awake infant fMRI is an exciting method that holds great potential, as evidenced by a growing body of findings across labs (7, 47). The results presented here from more than 750 scanning sessions and almost a decade of effort demonstrate the increasing feasibility of this method as well as what researchers entering the field should expect will impact data collection. Several factors such as age, sex, data collection practices, and experimental designs influence the success of a scan and should be carefully considered when designing and optimizing future experiments. Although any given scan is unpredictable, hopefully the present analyses might help other groups optimize future awake infant fMRI studies to advance understanding of the developing mind and brain.

## Acknowledgments

This work was supported by NSF Graduate Research Fellowship 2023350036 (LB), NIH grant R01HD103847 (RS), NIH grant R21HD090346 (RS), the Packard Fellowship (RS), the Canadian Institute for Advanced Research (NBTB), and the James S. McDonnell Foundation https://doi.org/10.37717/2020-1208 (NBTB).

We express deep appreciation to all of the families and infants who participated over the years for making this research possible. We also thank all members of our teams who contributed to the datasets summarized in this paper: at Yale, K. Armstrong, L. Rait, J. Daniels, and A. Letrou for recruitment, R. Watts for technical support, V. Bejjanki, L. Skalaban, A. Bracher and N. Córdova for implementation and data collection, and J. Fel and J. Cross for data collection and gaze coding; at MIT, K. Lydic for technical support, A. Fuchs, L. Herrera, I. Kawauchiya, J. Medrano, I. Nichoson, S. Riskin, S. Saba, B. Santi, M. Soza, and A. Zhou for data collection, A. Takahashi and S. Shannon at the Athinoula A. Martinos Imaging Center for MRI scanning and technical support, B. Keil for technical support, and K. Brewer from The Center for Brains, Minds, and Machines for photo and media support.

---

**Supplementary Figure 1:**
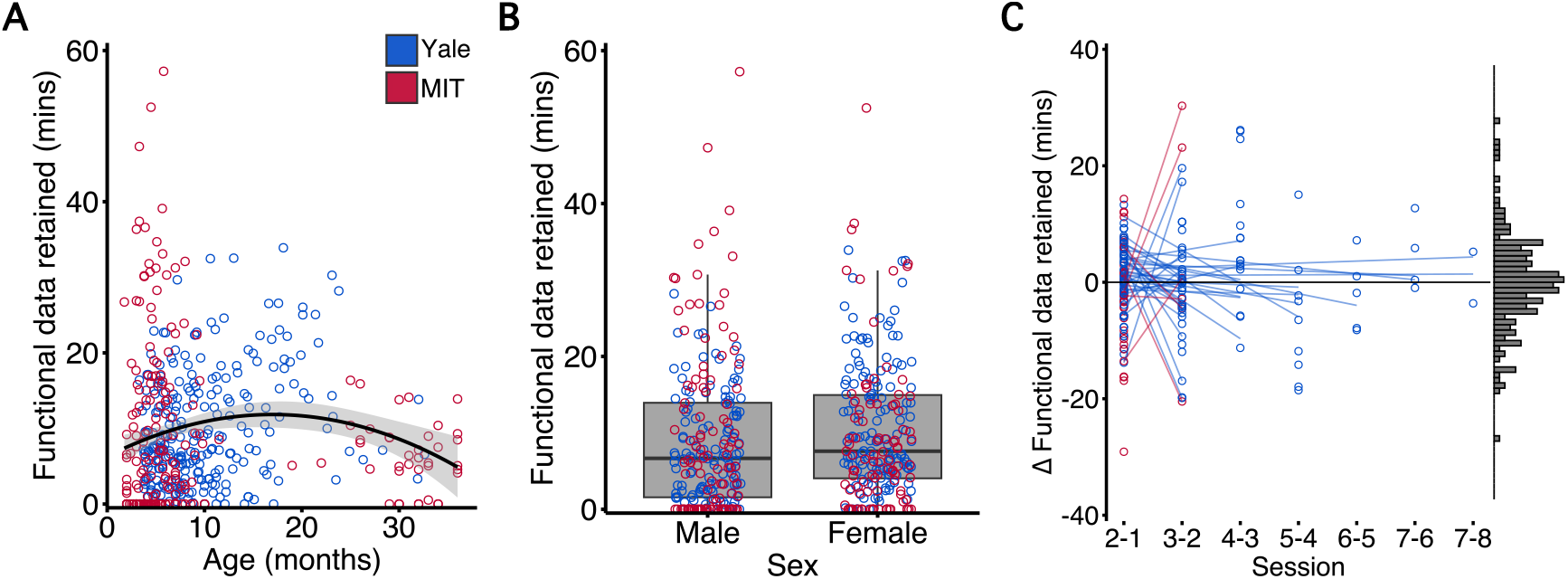
Predicting minutes of data retained with averaged MIT sessions. The analyses in the main text define a session as one visit to the scanning facility. At Yale, individual infants did not return again for at least a month. At MIT, infants were often invited back for multiple visits within a month and the aggregated dataset was treated as a session. Under the MIT definition (which does not impact Yale), data from 465 sessions (257 Yale, 208 MIT) where the infant entered the scanner and provided some amount of functional data were available to assess factors impacting the amount of usable data retained after motion exclusions (3.0-mm threshold at Yale, 0.5- or 1.0-mm thresholds at MIT). An average of 20.76 minutes (12.35 Yale, 31.15 MIT) of functional data were collected per session, of which 9.64 (46.4%) minutes were retained on average (10.04 Yale, 9.15 MIT; Range = [0 – 57.25], *Mdn* = 7.10, *IQR* = 11.97). A multiple regression model predicting the number of minutes of usable functional data retained per session as a linear function of institution, infant age, and assigned sex (hour of scan was not sensible for multi-visit sessions) did not provide a significant overall fit (*F*(3, 461) = 0.99, *p* = .398, *R*^2^ = .01, *R*^2^adj = .00). However, adding a linear+quadratic term for age — the superior model in the main text — yielded a significant fit (*F*(4, 460) = 3.22, *p* = .013, *R*^2^ = .03, *R*^2^adj = .02). The quadratic term for infant age was significant (*β* = -34.12, *p* = .002; 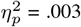), but sex (*β* = 1.01, *p* = .237; 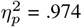) was not. Many of these infants (*N* = 97) attended multiple scanning sessions (range: 2–8; *N* = 270 sessions). Using the same approach from the main text to test for practice effects, the distribution of change scores (*M* = 0.00, *SD* = 9.09) again did not significantly differ from zero (*t*(172) = 0.00, *p* = .998). In a multiple regression model predicting the change in usable minutes, there was again no significant effect of session number (*β* = 0.56, *p* = .337) or infant age (*β* = -0.12, *p* = .304).

**Supplementary Figure 2:**
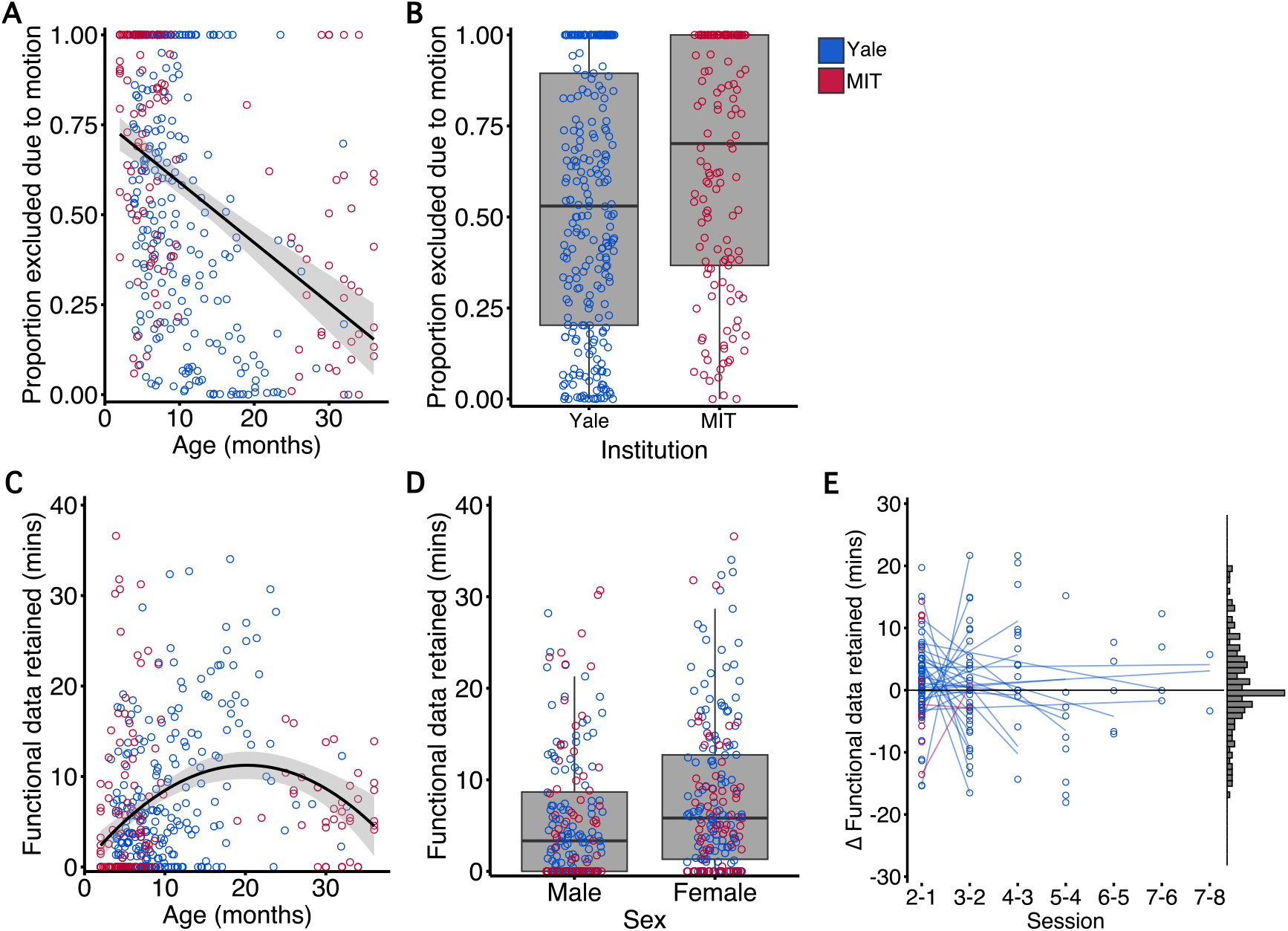
Applying MIT’s more lenient motion thresholds to Yale’s 10 – 18 month-olds. When harmonizing motion thresholds in the main text, the authors applied MIT’s 0.5 mm motion threshold (as they used for 1–10 months) to Yale’s data from 10 – 18 months (as well as for under 10 months). Here, MIT’s 1 mm motion threshold (as they used for 18–36 months) was applied to these participants instead. Data from 394 sessions (257 Yale, 137 MIT) were included in this analysis. In these sessions, an average of 14.58 minutes of functional data were collected, of which an average of 7.01 (48.1%) minutes were retained (Range = [0 – 36.60], *Mdn* = 4.53, *IQR* = 11.10). A multiple regression model predicting the proportion of data excluded from a session due to head motion as a function of institution, infant age, assigned sex, and hour of scan provided a significant overall fit (*F*(4, 389) = 27.50, *p* < .001, *R*^2^ = .22, *R*^2^adj = .21). In addition to age (*β* = -0.02, *p* < .001; 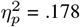) and institution (*β* = 0.14, *p* < .001; 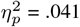), assigned sex was also a significant predictor (*β* = -0.08, *p* = .014; 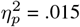). These results align with what was found with the other thresholding approach reported in the main text and shown in Figure 4. An additional multiple regression model predicting the number of minutes of usable functional data retained from each session as a function of institution, linear+quadratic infant age, assigned sex, and hour of scan provided a significant overall fit (*F*(5, 388) = 9.76, *p* < .001, *R*^2^ = .11, *R*^2^adj = .10). The quadratic term for infant age (*β* = -45.52, *p* < .001; 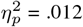), assigned sex (*β* = 1.69, *p* = .025; 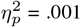), and the linear term for infant age (*β* = 22.47, *p* = .003; 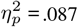) were significant predictors; this matches what was found with the other thresholding approach reported in the main text and shown in Figure 5. For participants with multiple sessions, the distribution of change scores (*M* = 0.97, *SD* = 8.15) did not differ significantly from zero (*t*(150) = 1.46, *p* = .148). In a multiple regression model predicting the change in usable minutes, there was again no significant effect of session number (*β* = 0.24, *p* = .651) or infant age (*β* = -0.11, *p* = .314).

**Supplementary Figure 3:**
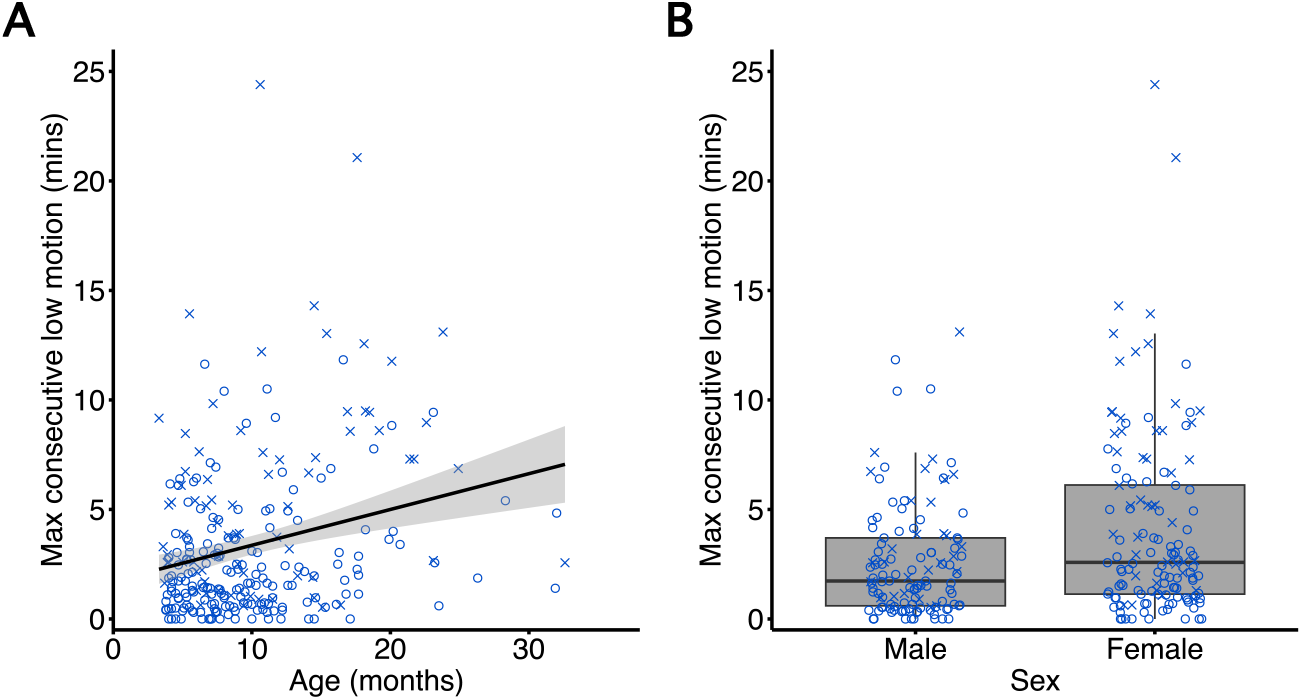
Predicting consecutive minutes of low motion data obtained. Certain study designs and analysis types benefit from continuous timecourses of fMRI data (i.e., periods during which no frames are removed due to head motion above Yale’s 3.0 mm threshold). From each session during which functional data were collected in Yale (*N* = 257), the authors calculated the longest span of low-motion data (*M* = 3.38, *SD* = 3.57 min). A multiple regression model predicting this span as a linear function of infant age, assigned sex, and hour of scan provided a significant overall fit (*F*(3, 253) = 9.53, *p* < .001, *R*^2^ = .10, *R*^2^adj = .09). Infant age (*β* = 0.15, *p* < .001; 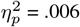) and sex (*β* = 1.34, *p* = .002; 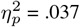) were significant predictors. Post-hoc analyses were conducted to verify the linearity of the age effect. BIC for the model with a linear age term (1,383.14) was lower than for a model with linear and quadratic age terms (1,753.49), suggesting that the linear model better balanced fit and complexity. In many cases (N = 74), the longest span of low-motion data was terminated by the researcher (i.e., at the end of an experiment) and not because of infant motion. These sessions are indicated in the figure with an “x”.

## Notes

### Competing Interest Statement

The authors have declared no competing interest.

### Summary of Updates

We revised the title of the manuscript and amended the Introduction to better convey the rationale for the study. We conducted a series of new analyses reported in the Results, Figures, and Supplement to harmonize the motion thresholds used during preprocessing across labs. We visualized the data collection apparatus used by each lab in a figure so that readers can more easily understand and compare approaches. We reported the number and demographics of infants included at each stage of the analysis pipeline in a new figure. Finally, we expanded the Discussion to provide a deeper synthesis of the differences and relative strengths and weaknesses between approaches.

